# Super-Resolution Tracking of Mitochondrial Dynamics with An Iridium(III) Luminophore

**DOI:** 10.1101/312371

**Authors:** Qixin Chen, Chengzhi Jin, Xintian Shao, Ruilin Guan, Zhiqi Tian, Chenran Wang, Fei Liu, Peixue Ling, Jun-Lin Guan, Liangnian Ji, Fengshan Wang, Hui Chao, Jiajie Diao

## Abstract

Combining luminescent transition metal complex (LTMC) with super-resolution microscopy is an excellent strategy for the long-term visualization of the dynamics of subcellular structures in living cells. However, it remains unclear whether iridium(III) complexes are applicable for a particular type of super-resolution technique, structured illumination microscopy (SIM), to image subcellular structures.

As described herein, we developed an iridium(III) dye, to track mitochondrial dynamics in living cells under SIM. The dye demonstrated excellent specificity and photostability and satisfactory cell permeability. While using SIM to image mitochondria, we achieved an approximately 80-nm resolution that allowed the clear observation of the structure of mitochondrial cristae. We used the dye to monitor and quantify mitochondrial dynamics relative to lysosomes, including fusion involved in mitophagy, and newly discovered mitochondria-lysosome contact (MLC) under different conditions. MLC remained intact and fusion vanished when five receptors, p62, NDP52, OPTN, NBR1, and TAX1BP1, were knocked out, suggesting that these two processes are independence.

## Introduction

Highly mobile and dynamic in living cells, mitochondria are the energy-generating organelles of cell^1, 2, 3^, the disorder of which is associated with various diseases, including Alzheimer’s, Parkinson’s, and cancer^4, 5^. Recently developed super-resolution fluorescence microscopies such as stimulated emission deletion (STED)^6, 7, 8^, structured illumination microscopy (SIM)^9, 10, 11^ and stochastic optical reconstruction microscopy (STORM)^12, 13, 14^, as well as other single-molecule super-resolution imaging techniques^15, 16, 17^, are enhanced new tools for investigating the dynamics of subcellular structures, including mitochondria. At the same time, they have also added to the requirements of fluorescent dyes, which need especially low cytotoxicity and high photostability to make imaging living cells possible.

Currently, imaging subcellular structures relies upon fluorescent proteins^18, 19^, organic dyes^20^, and quantum dots^21^; however, none of those dyes are suitable for tracking the dynamics of subcellular structures due to their poor photostability and vulnerability to photobleaching^22^. Studies have shown that using luminescent transition metal complexes (LTMCs)^23, 24, 25, 26^, including Ru, Re, Pt, Au, and Zn, is an excellent alternative strategy that can overcome those drawbacks. For example, to image mitochondria, Tang et al. reported a Zn(II) complex dye whose fluorescence intensity decayed to 10% after a short period of continuous scan under STORM^27^. More recently, the lab of Jim A. Thomas developed a Ru(II) complex dye with extreme photostability, large Stokes shift and subcellular targeting to image nuclear chromatin and mitochondria at a resolution of less than 50 nm under STED^28^. However, it remains unclear whether third-row LTMC dyes are applicable with SIM to image the dynamics of subcellular structures.

Among LTMC dyes, iridium(III) complexes are the most attractive candidate for bioimaging applications due to their high phosphorescent quantum yield at room temperature, high penetration and emission spectrum that can be extended to near-infrared areas, excellent photochemical and physicochemical stability that allows for long-term imaging, high biological safety, and low cytotoxicity^4, 29^. Those unique structures and photophysical properties make it possible to develop innovative bioimaging applications based on phosphorescence.

As described herein, we developed a small molecule dye based on iridium(III) complex for tracking mitochondrial dynamics under a SIM for the first time. The dye not only has exceptional cell permeability, light stability and mitochondrial specificity but can also allow the observation of mitochondria at up to 80-nm resolution in living cells under SIM. Moreover, the dye permits the clear observation of the structure of mitochondrial cristae, as well as the recording of dynamic approach and separation processes of mitochondria. We applied the dye to monitor fusion involved in mitophagy and mitochondria–lysosome contact (MLC) in living cells and found that five receptors, p62, NDP52, OPTN, NBR1, and TAX1BP1, for mitophagy played no role in regulating MLC upon stimulation. Our findings thus illustrate a novel perspective for using LTMC dyes such as iridium(III) complexes to image the dynamic processes of subcellular structures in living cells under SIM.

## Results

### Synthesis and optical characterization

First, we designed and synthesized an iridium(III) complex dye with a molecular weight of 970 Da to specifically image mitochondria in living cells under SIM (Fig. 1a). The main synthesis reaction route consisted of three major steps (Fig. 1b), and the characterizations with electrospray mass spectrometry and nuclear magnetic resonance spectroscopy appear in Supplementary Fig. 1-3. The dye had higher phosphorescence at a wavelength of approximately 700 nm and different optical phosphorescence characteristics in phosphate-buffered saline at pH 4-10 (Fig. 1c-d), which indicates that it can be used in living cells in different pH environments. The emission spectra show that the luminescent intensity in pH 4-6 is relatively low compared to that in pH 6-10. This phenomenon may contribute to the presence of pyrazine ring in the auxiliary ligands of iridium(III) complexes. Since the phosphorescence intensity of this iridium(III) complex is relatively stable in the pH range of 6.0 - 10.0, it is acceptable for mitochondrial imaging (physiological pH: 6.50 to 8.20)^30^. To further investigate the optical properties of the dye in other solvent media, we performed fluorescence detection in 100% fetal calf serum (FBS) and Dulbecco’s modified eagle medium (DMEM; Supplementary Fig. 4a-b). The dye exhibited high fluorescence intensity in FBS and low fluorescence in DMEM, which indicates that it can obtain a lower background in DMEM under SIM. Upon testing the phosphorescence properties of the dye at different temperatures, we found that it had the same optical properties at all temperatures tested (Supplementary Fig. 4 c-d), which suggests using the dye can afford consistent data acquisition at physiological temperature (37 °C) and ambient room temperature (25 °C) for SIM detection in living cells.

**Figure 1.**
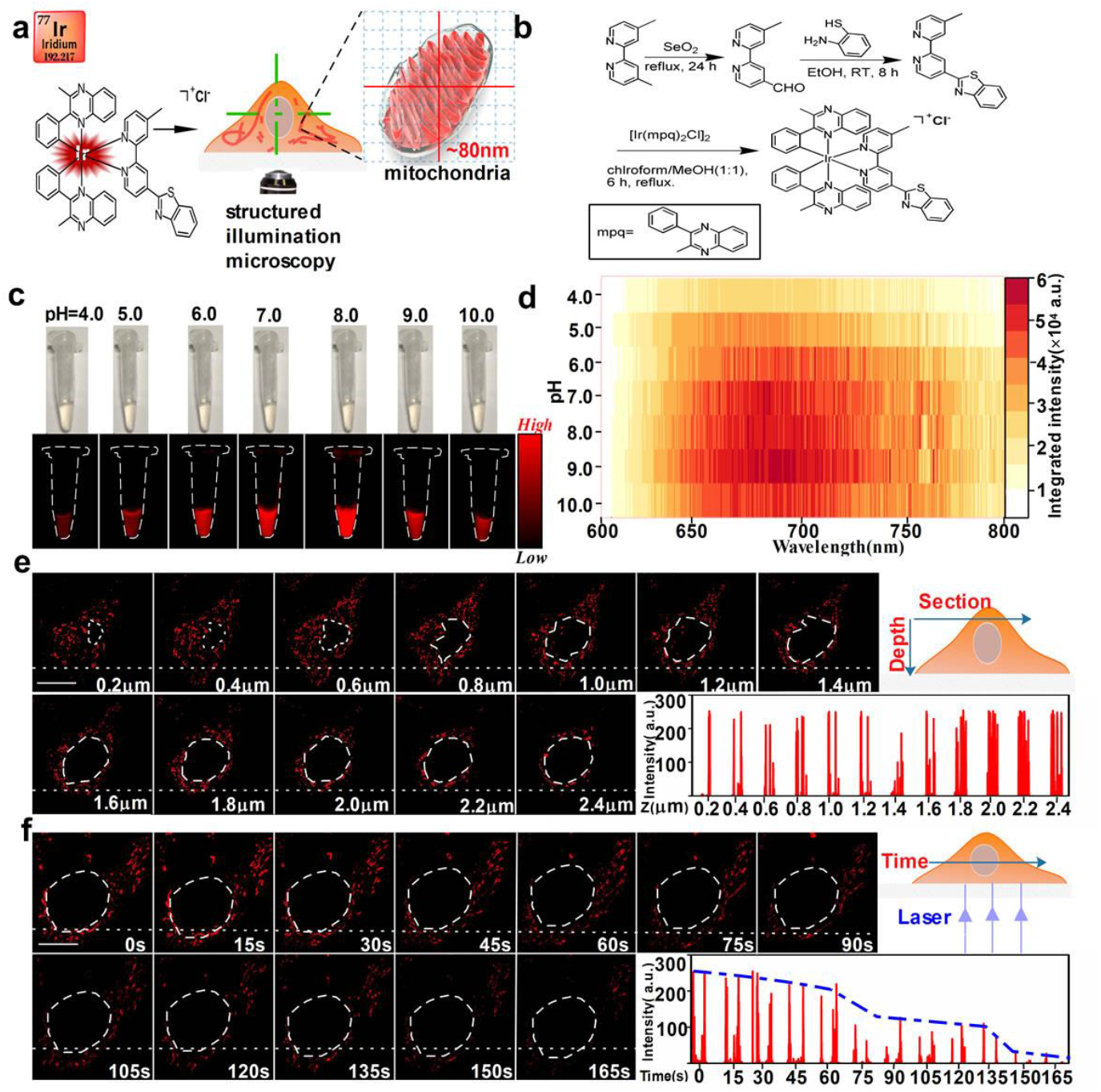
Synthesis and optical characterization of the iridium(III) complex dye. **(a)** Schematic representation of the iridium(III) complex dye while imaging mitochondria under SIM. **(b)** Synthesis route of the iridium(III) complex dye. **(c)** Fluorescence image of the iridium(III) complex dye in phosphate-buffered saline at different pH levels. **(d)** Photoluminescence mapping of the iridium(III) complex dye. **(e)** Mitochondrial images at different depths using the iridium(III) complex dye. White dotted lines show fluorescence intensity of a single mitochondrion image at every depth of 0.2 μm. **(f)** Photobleaching properties of the iridium(III) complex dye under laser stimulation for 165 s continuously recorded in living cells. SIM frames were deliberately spaced at 15-s intervals. Scale bars: **(e)** 5 μm, **(f)** 5 μm.

### Characterization of the iridium(III) complex dye in living cells

To investigate the cell permeability of the dye in living cells, concentrations of dye ranging from 0.1 to 50 μM were incubated with HeLa cells for 30 min prior to observation under a fluorescence microscope.

We observed high phosphorescence accumulation, and even nuclei were stained when cells were exposed to dye at concentrations of 5-50 μM, whereas at concentrations of 0.1, 0.25, and 0.5 μM, phosphorescence intensity was too low to be visible (Supplementary Fig. 5). Taking both into consideration, we determined that the dye at concentrations of 1 and 2.5 μM could be used to image mitochondria well.

Next, we used flow cytometry to quantitatively analyze the cellular uptake of the dye (Supplementary Fig. 6). Results indicated that when the cells were exposed to dye at concentrations greater than 5.0 μM, the cells took up excessive dye, whereas at a concentration less than 2.5 μM, the cellular uptake of the dye gradually decreased in a dose-dependent manner. Thereafter, we evaluated the cytotoxicity of the dye at concentrations of 0.1-50 μM during different durations of incubation using a CCK-8 assay (Supplementary Fig. 7). Under continuous 12-hour co-culture, results revealed no significant differences in cell viability between the control group and the groups treated with the dye at concentrations of less than 10 μM.

Although it is feasible to observe and record dynamic progresses at the cellular level with confocal optical fluorescence microscopy, it is difficult to distinguish subcellular structures at resolutions less than 200 nm due to the Abbe diffraction limit^4, 31, 32^. In response to that problem, after clarifying our dye’s cell permeability, high specificity, and low cytotoxicity, we investigated differences between the dye imaging of mitochondria under confocal microscopy and SIM (Supplementary Fig. 8, Supplementary Movies 1 & 2). To avoid the potential cytotoxicity of high-concentration dyes, we selected 0.5, 1, 2.5, and 5 μM as experimental concentrations. Results revealed that the dye had a low phosphorescence background in confocal microscopy and SIM, as well as that the resolution of the dye at a 1 μM concentration was superior to those at 0.5, 2.5, and 0.5 μM concentrations (Supplementary Fig. 8c, f, i, l). When concentrations exceeded 1 μM, a large amount of the dye was taken up by the cells, which caused strong phosphorescence (Supplementary Fig. 8a-f) that did not meet the requirement for imaging mitochondria at the nanoscale level. By contrast, when the concentration was less than 1 μM, less dye was taken up by the cells, which resulted in weak phosphorescence due to which not even mitochondrial morphology was visible (Supplementary Fig. 8i, j). Therefore, we chose 1 μM as an optimal concentration for further study.

To investigate the photostability and penetration depth of the dye at 1 μM for imaging mitochondria in living cells under SIM, we recorded a single mitochondrion image at every 0.2 μm of depth (Fig. 1e). Results indicated that the dye had uniform tissue permeability at different depths in living cells. The photobleaching properties of the dye directly affected the monitoring of mitochondrial dynamic processes^33, 34^. Next, we performed a long-term continuous laser (165 s) to stimulate cells in order to monitor the photostability of the dye in the same section of the cell (Fig. 1f). We found that the dye had high phosphorescence within 60 s of continuous laser stimulation and could image mitochondria within 135 s without photobleaching. Those properties allowed us to record more dynamic information while monitoring mitochondria in living cells under SIM with the dye.

### Whole-cell 3D SIM images of mitochondrial ultrastructures using the iridium(III) complex dye

Visualizing mitochondrial ultrastructures affords new understandings of the pathology and diagnosis of mitochondria-related diseases^35, 36^. To investigate the capability of obtaining more information about mitochondrial ultrastructures, we used the dye to image mitochondrial ultrastructures in living cells under 3D SIM. Shown in Fig. 2a, results revealed that after cell incubation with the dye for 30 min, intracellular mitochondria showed spherical, rod-shaped, or filamentous particles approximately 2.0 μm long (Fig. 2c-2) and approximately 0.7 μm wide (Fig. 2c–3), which is consistent with normal mitochondrial volume in HeLa cells (width: 0.5-1.0 μm, length: 1.5-3.0 μm). The lamellar cristae in mitochondrial were also visible using the dye—cristae thickness was approximately 105 nm (Fig. 2b, Supplementary Fig. 9)—and the dye could be evenly distributed on mitochondria (Fig. 2d, Supplementary Movie 3), which suggests that it can be located on the mitochondrial membrane and have high specificity. To further clarify the resolution of the dye for imaging mitochondria, we obtained a full-width at the half-maximum (FWHM) up to 80 nm under SIM (Fig. 2e-f), which is similar to the Atto 647 mitochondrial dye reported by Han et al. (FWHM: 91 nm)^34^. Such results suggest that our dye offers higher resolution and precision for tracking mitochondria under SIM.

**Figure 2.**
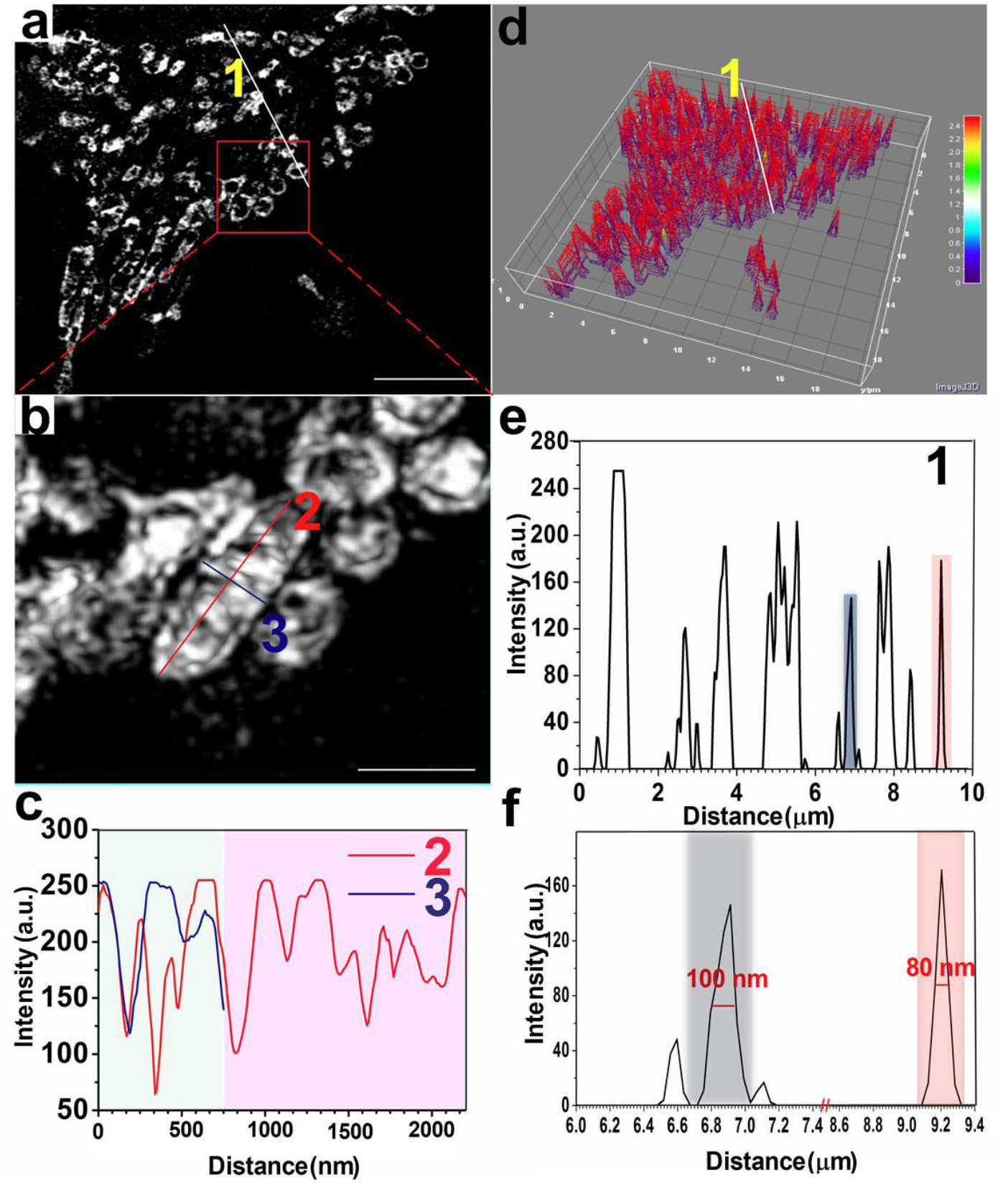
Iridium(III) complex dye images of mitochondrial ultrastructure. **(a)** Iridium(III) complex dye images of mitochondria in 1 μM concentration under 3D SIM. White solid line **1** shows fluorescence intensity, and red rectangle shows partial amplification. **(b)** Single mitochondrial local enlargement of **(a)**. Red solid line **2** shows the length of a single mitochondrion, and blue solid line **3** shows its width. **(c)** Individual mitochondria fluorescence intensity with length as red solid line **2** and width as blue solid line **3. (d)** iridium(III) complex dye 3D map distributed in mitochondria under SIM. **(e)** Fluorescent intensity distribution of white solid line **1. (f)** Local magnification of **(e)**, with resolution of iridium(III) complex dye image of mitochondria up to 80 nm. Scale bars: **(a)** 5 μm, **(b)** 1 μm.

### Iridium(III) complex dye for tracking mitochondrial dynamics

Mitochondria rank among the most dynamic organelles in cells^34^, and understanding their dynamic processes is important for analyzing the causes of many diseases^37^. Therefore, a strategy for visualizing the dynamics of mitochondria at nanoscale has important implications for understanding mitochondria-related diseases. With that goal in mind, we recorded the process of mitochondrial dynamics using our dye after incubating it in live cells for 30 min (Fig. 3, Supplementary Movie 4). We observed that two mitochondria (Fig. 3d-f, red arrow) maintained a distance of approximately 0.6 μm in Frame 1 (Fig. 3b-1) and gradually approached each other in Frames 1 and 2 (Fig. 3b, e, f). We also observed the separation process of mitochondria, particularly the gradual disintegration from an intact mitochondria (Fig. 3c, g-i, blue arrows), which suggests that our dye can be used to study the dynamic changes such as fusion and fission of mitochondria.

**Figure 3.**
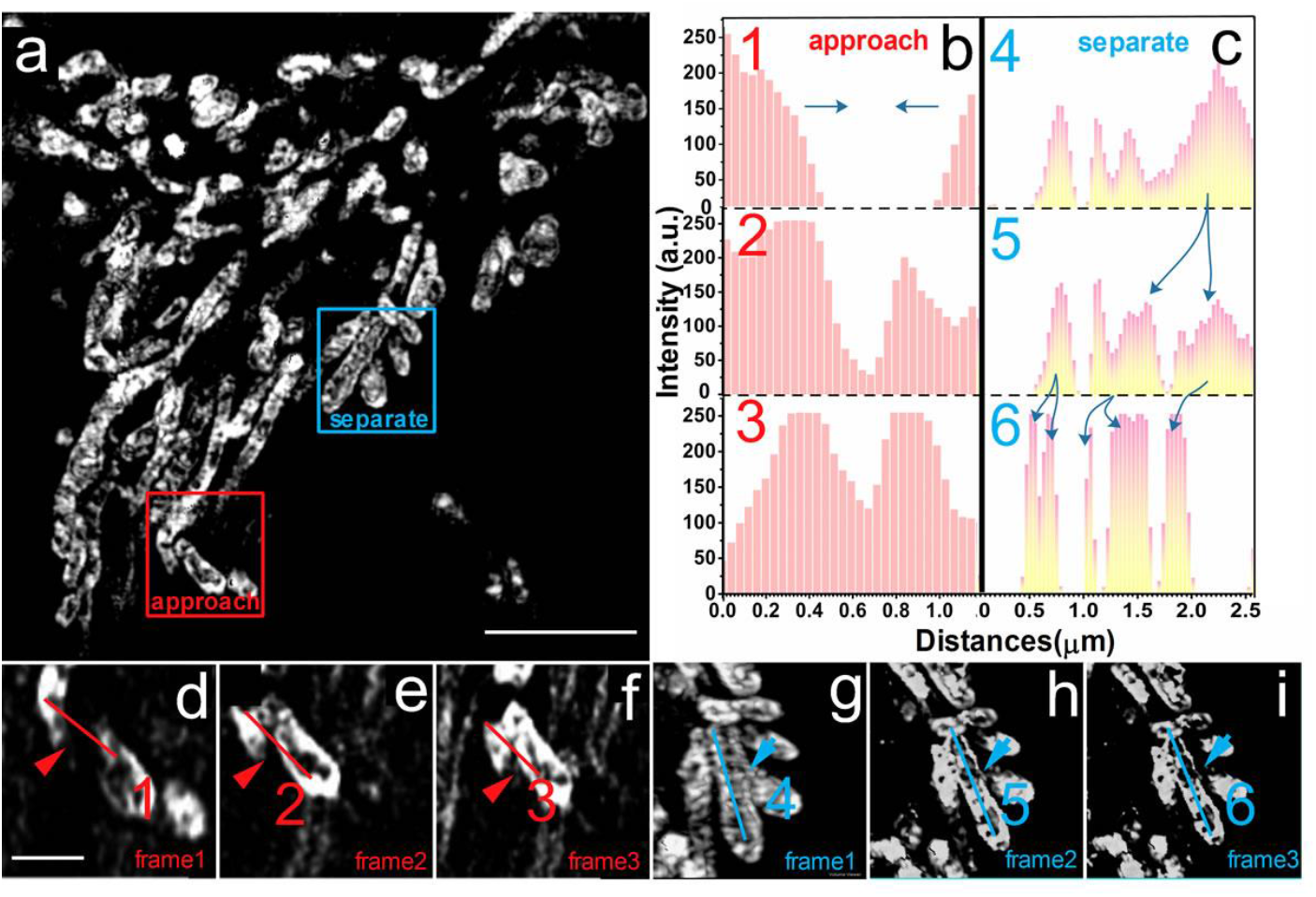
Iridium(III) complex dye tracking of mitochondrial dynamics under SIM. **(a)** Iridium(III) complex dye images of mitochondria under SIM. Red rectangles indicate dynamic processes of approach, and blue rectangles indicate dynamic processes of separation. **(b)** Trend of fluorescence intensity in the dynamic process of approach. **(c)** Trend of fluorescence intensity in the dynamic process of separation. **(d-f)** Frames 1-3 of the dynamic approach process in the red rectangle of **(a). (g-i)** Frames 4-6 of the dynamic separation process in the blue rectangle of **(a)**. The time interval between each frame is approximately 2 s. Both red and blue solid lines show fluorescence intensity. Scale bars: **(a)** 5 μm, **(d)** 1 μm.

### Application of the iridium(III) complex dye to track MLC and fusion

The interactions of mitochondria and lysosomes, including their fusion involved in mitophagy, are essential for repairing damaged mitochondria. Recently, a direct contact between mitochondria and lysosome was demonstrated in normal living cells^38^. To investigate the crosstalk of mitochondria and lysosomes, we used our dye together with commercial LysoTracker Green to label lysosomes in mammalian cells. We confirmed that mitochondria and lysosomes were close to each other to form MLC, similar to what has been previously reported^38^. We observed that MLC events were normal in wild-type (WT) cells (Fig. 4a-f, Supplementary Movie 5-7), but did not observe fusion. Moreover, we found two types of MLCs (Fig. 4c), point contact with limited overlap (Fig. 4d) and extended contact showing an elongated contact surface (Fig. 4e).

**Figure 4.**
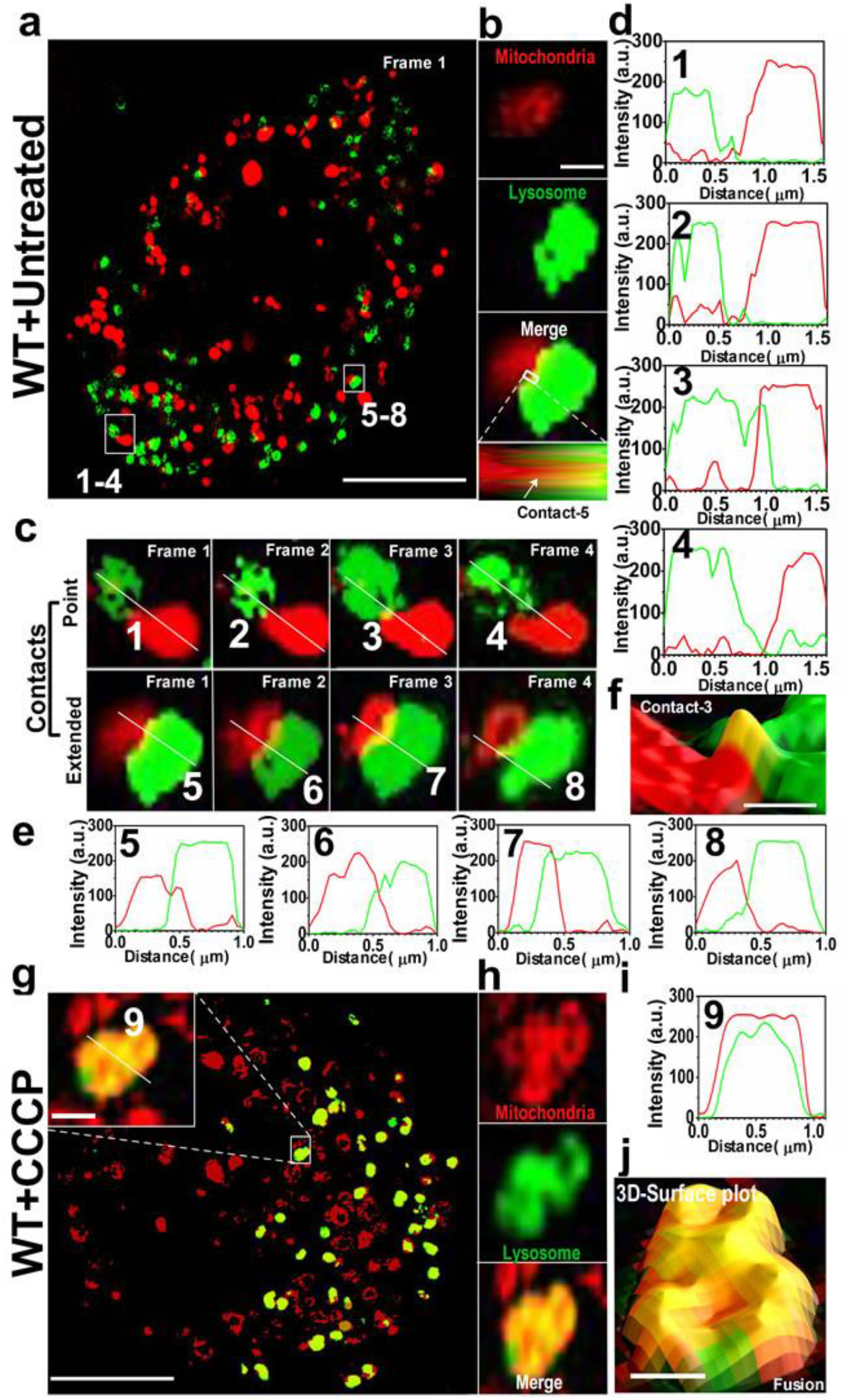
Iridium(III) complex dye tracking of mitochondria-lysosome contact (MLC) and fusion events in living cells. **(a)** MLC events in WT cells. White rectangles represent two types of MLCs, point (1-4) and extended (5-8) contacts. **(b)** A representative MLC event. White rectangle represents the amplification. **(c)** Point and extended MLCs in **(a)**. White solid lines indicate the fluorescence intensity shown in **(d)** and **(e). (f)** Partial enlargement of **(d)**-3. **(g)** Fusion events in WT cells after treatment CCCP. White rectangle shows the representative fusion event, and the white solid line shows fluorescence intensity. **(h)** A representative fusion event detected by using the iridium(III) complex dye (red, mitochondria) and LysoTracker Green (green, lysosome). **(i)** Fluorescence intensity of the solid line of **(g)**-9 inset. **(j)** 3D SIM surface plot of merged image in **(h)**. Scale bars: **(a)** 5.0 μm, **(b)** 0.5 μm, **(e)** 0.2 μm, **(g)** 5.0 μm, **(g)**-9 0.5 μm, **(j)** 0.5 μm.

For the application of our dye to image damaged mitochondria related to pathological conditions, we performed another experiment with inducer treatment. We used 10.0 μM carbonyl cyanide m-chlorophenyl hydrazone (CCCP), a common mitochondria damage inducer, to treat cells 12 h prior to staining with our dye and LysoTracker Green. Compared to untreated cells (Fig. 4g-j, Supplementary Movie 8), we observed a significant increase of large overlaps (yellow spots) of mitochondria and lysosomes corresponding to the fusion after CCCP treatment.

### Iridium(III) complex dye to track MLC in Penta knockout HeLa cells

MLCs were observed in both normal and stimulated conditions, while fusion involved in mitophagy was only found after CCCP treatment. Pathologically, PINK1 is recruited to the mitochondrial membrane to phosphorylate Ser65 of ubiquitin ligase and trigger mitophagy. During that process, several receptors (i.e., p62, NDP52, OPTN, NBR1, and TAX1BP1) play an important role^39^. Therefore, it is reasonable to check whether these mitophagy proteins are involved in MLC events other than the recently reported RAB7GTP hydrolysis^38^. To investigate the correlation between MLC and fusion, we performed the SIM experiment in Penta knockout (KO) HeLa cells which are deficient of mitophagy. After using 10 μM CCCP to treat cells and staining with our dye and LysoTracker Green, we found MLC events including 2-3 faint yellow spots for point MLC throughout the cells (Fig. 5a-c, Supplementary Movie 9). In addition, we recorded the evolution of an MLC event (Fig. 5d-e, Supplementary Movie 10), in which a mitochondrion and a lysosome underwent approach, contact, and separation. Meanwhile, fusion event involved in mitophagy in Penta KO HeLa cells (Fig. 5a) was disappeared compared to what was found in the WT HeLa cells (Fig. 4g). To that end, we performed a controlled experiment in which Penta KO HeLa cells did not receive CCCP treatment. We detected no significant change of MLC events in Penta KO cells (Fig. 5g, Supplementary Movie 11), which indicates that fusion between mitochondria and lysosomes are independent to MLC. Again, our dye can be used to monitor MLC events in living cells.

**Figure 5.**
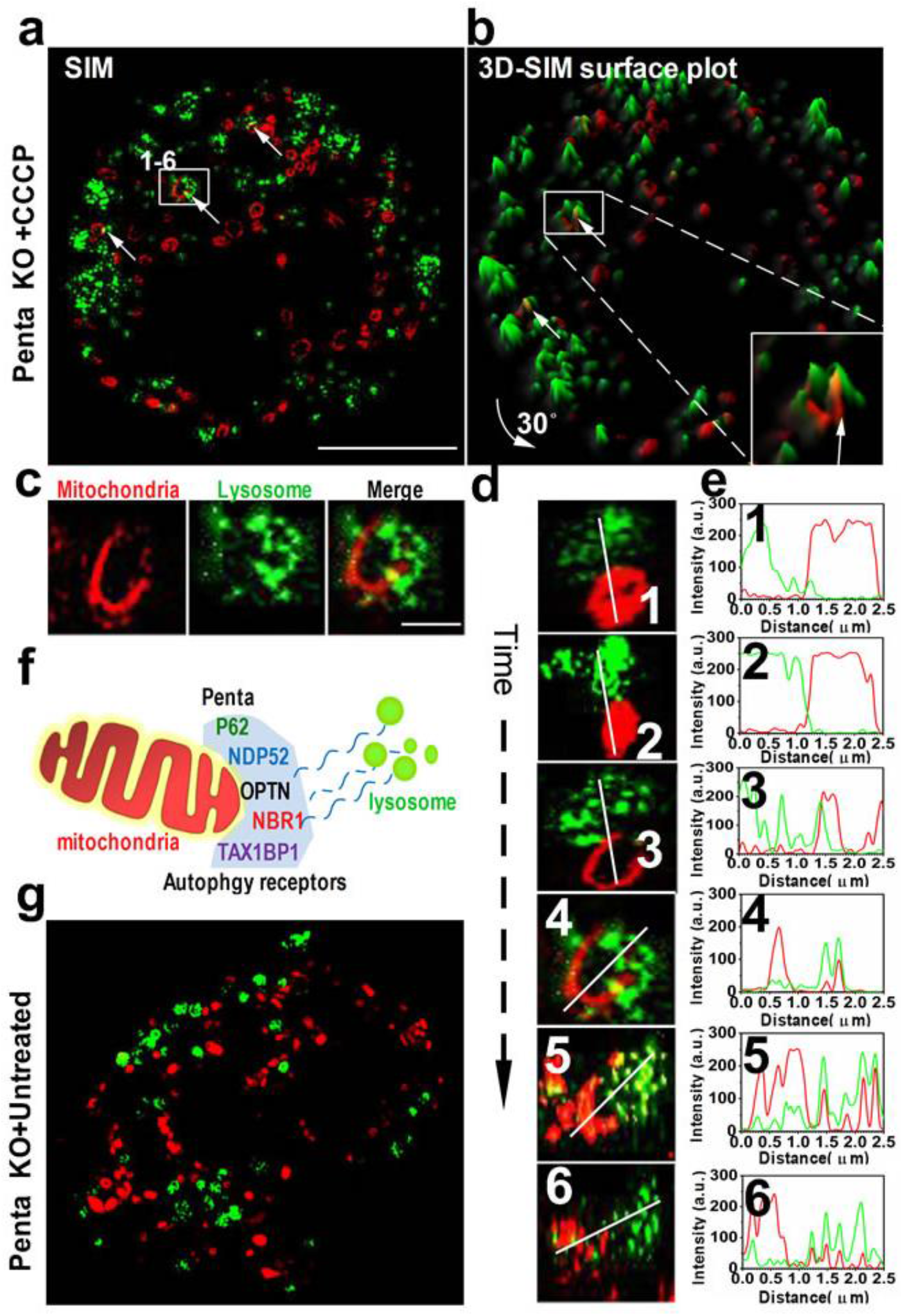
Iridium(III) complex dye for tracking MLC in Penta knockout HeLa cells. **(a)** MLC events in Penta KO cells treated with 10 μM CCCP for 12 h. White rectangle represents the MLC shown in **(d). (b)** 3D SIM surface plot of **(a)** after 30-degree rotation. White arrows indicate the MLC events. **(c)** A representative MLC event. **(d)** Time evolution of one MLC in living cell. White solid lines indicate fluorescence intensity shown in **(e)**. Scale bars: **(a)** 5.0 μm, **(b)** 1.0 μm.

### Quantitative analysis of the interaction between mitochondria and lysosomes

To quantitatively assess the distance changes between mitochondria and lysosome under different conditions, we propose a Di-value driven from FWHM by the calculation formula shown in Fig. 6a. FWHM refers to the full width of the image at half-maximum value and is a direct indicator of the resolution. We then apply the Di-value to quantify the distance change between mitochondria and lysosomes under different conditions. We first calculated the *D*i value (0.884 ± 0.116, n = 10) in untreated WT cells for MLC events. We then analyzed the *D*i value of WT cells treated with CCCP. We found a much lower Di value (0.146 ± 0.118, n = 10) for fusion events, indicating a significant difference upon stimulation (Fig. 6b). For Penta KO cells with or without CCCP, no significant difference in Di values was observed (1.123 ± 0.176 *vs* 1.128 ± 0.140, n = 10) (Fig. 6c).

**Figure 6.**
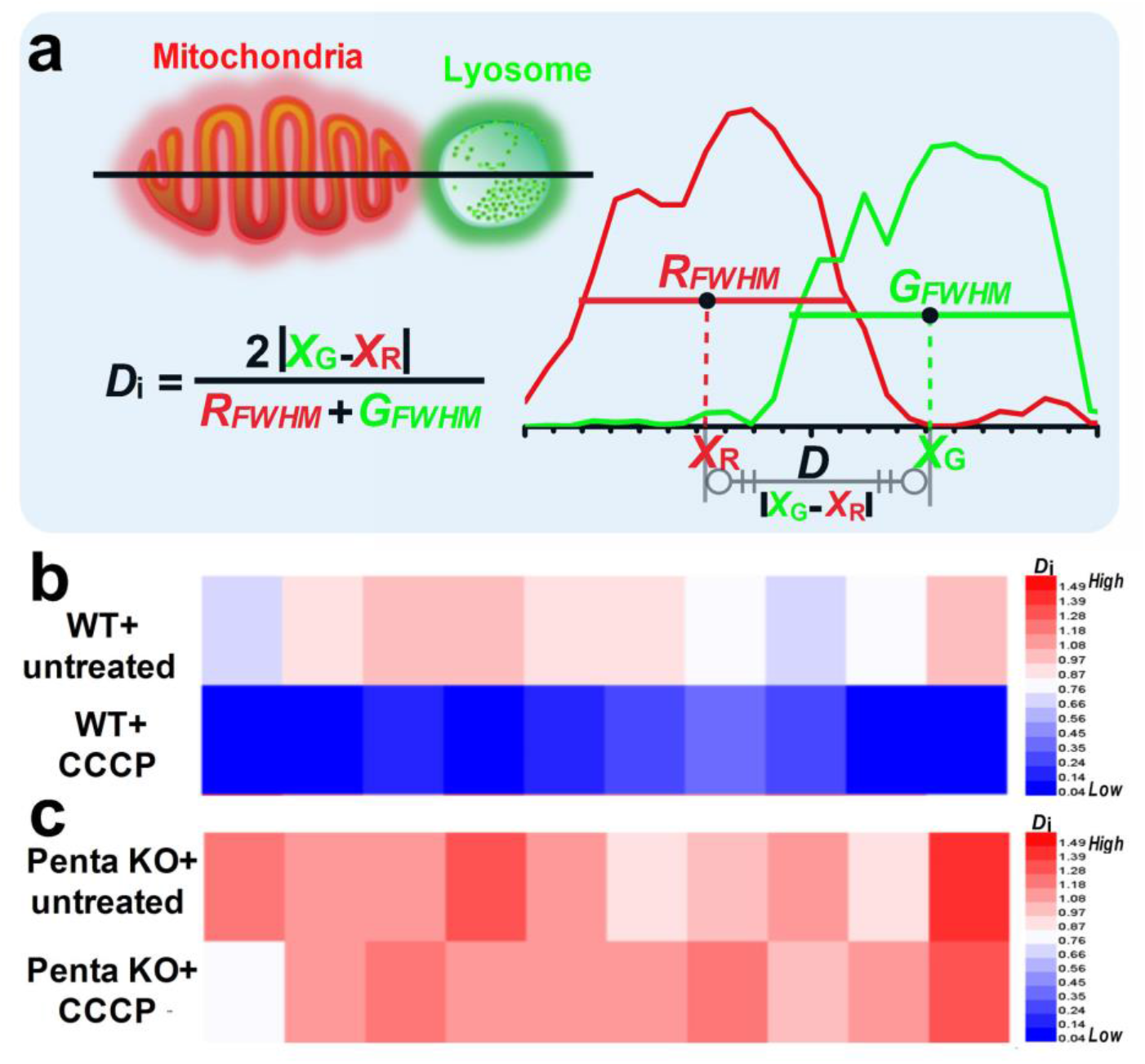
Quantitative analysis of the interaction between mitochondria and lysosomes. **(a)** The schematic illustration of a *D*i-value for quantitative analysis of the distance between mitochondria and lysosomes. *R*_FWHM_ indicates the FWHM of red color (mitochondria); *G*_FWHM_ indicates the FWHM of green color (lysosomes); *X*_R_ indicates the X-axis value of the *R*_FWHM_ center; and *X_G_* indicates the X-axis value of the *G*_FWHM_ center. **(b)** *D*i-values of mitochondria and lysosome in WT cells with or without CCCP treatment (n = 10). **(c)** Di values of mitochondria and lysosome in Penta KO cells with or without CCCP treatment (n = 10).

## Discussion

We discovered that a third-row transition metal complex fluorescent dye based on iridium(III) can be used to track mitochondrial dynamics under SIM. Using our dye, we obtained a mitochondrial image with approximately 80-nm resolution. We observed mitochondrial cristae and recorded the dynamic approach and separation of mitochondria. Our results provide a novel perspective on using LTMC dyes such as an iridium(III) complex to image the dynamic process of subcellular structures in living cells under SIM. Given the synthesis of the iridium(III) complex, future work could focus on developing a variety of novel iridium(III) dyes for the specific imaging of lysosomes, mitochondria, cell membranes, nuclei, and other organelles under SIM toward the eventual goal of mapping the organelle interaction network and thereby clarifying the network’s establishment, maintenance, dynamic changes, and regulatory mechanisms and to reveal its physiological and pathological functions.

Mitochondrial dynamic processes including fusion and fission are closely related to many diseases^40^. With our iridium(III) complex dye, we have demonstrated the capability of imaging these dynamics with a high special resolution. Moreover, this new iridium(III) complex dye can also be used to quantitatively study the functional crosstalk between mitochondria and lysosomes, such as the fusion and MLC. The fusion between mitochondria and lysosomes is an important step of mitophagy for recycling damaged mitochondria^41^. We observed that the fusion of mitochondria and lysosomes can be significantly enhanced upon mitochondrial damage. However, the recently reported formation of MLCs differed from these fusion events of mitophagy in which mitochondria targeted to lysosomes for destruction^38^. Five receptors (i.e., p62, NDP52, OPTN, NBR1, and TAX1BP1) that play essential roles in mitophagy were not involved in the formation of MLCs^38^, which suggests MLCs and mitophagy are independent processes.

## Methods

### Materials

IrCl_3_·xH_2_O were purchased from Alfa Aesar, 4,4′-dimethyl-2,2′-bipyridine, diatomite, selenium dioxide 1-phenyl-1,2-propane-dione, 2-aminobenzenethiol and benzene-1,2-diamine were purchased from J&K Scientific. Lyso Tracker Green were purchased from Invitrogen (Invitrogen, Eugene, OR,USA); cell counting kit-8 (CCK-8) was obtained Dojindo Laboratories (Dojindo Laboratories, Kumamoto, Japan); fetal bovine serum (FBS), Dulbecco’s modified Eagle’s medium (DMEM), and other cell culture reagents were obtained from Gibco BRL (Grand Island, NY, USA).

### Synthesis and characterization of iridium(III) complex dye

The dye was synthesized by using a previously reported protocol^2^. 4′-methyl-[2,2′-bipyridine]-4-carbaldehyde and the auxiliary ligand 2-methyl-3-phenylquinoxaline (mpq)^43^ were synthesized according to literature methods. The iridium(III) dimmer [Ir(mpq)_2_Cl]_2_ was synthesized by using the similar method of [Ir(ppy)_2_Cl]_2_^44^. The synthetic process of the main ligand and the iridium(III) complex were according to our previous work.

The main ligand (2-(4′-methyl-[2,2′-bipyridin]-4-yl)benzo[d]thiazole (mbbt)) was synthesized by slowly dropping 2-aminobenzenethiol (196 mg, 1.55 mmol) into the EtOH solution of 4′-methyl-[2,2′-bipyridine]-4-carbaldehyde (297 mg, 1.5 mmol). After stirring overnight in RT, the product was condensed and then recrystallized by using CH_2_Cl_2_/ethanol to get yellow flaky solid Yield, 80.5%, 367 mg. Anal. Calcd. for C_18_H_13_N_3_S (%): C, 71.26; H, 4.32; N, 13.85. Found (%): C, 71.03; H, 4.63; N, 13.61.^1^H NMR (500 MHz, CDCl_3_) δ 9.27 (s, 1H), 8.60 (d, *J* = 1.0 Hz, 2H), 8.42 (s, 1H), 8.11 (s, 1H), 8.00 (d, *J* = 4.5 Hz, 2H), 7.49 – 7.45 (m, 2H), 7.20 (s, 1H), 2.54 (s, 3H). ES-MS, (CH_3_OH): m/z = 304.15 [M+H]^+^).

The goal iridium complex were synthesized by mixing mbbt (30.4 mg, 0.1 mmol) and [Ir(mpq)2Cl]2 (66.3 mg, 0.0525 mmol) in a degassing mixture of chloroform and MeOH (1:1, 40 ml). Then the solution was refluxed overnight in argon atmosphere. After the reaction was stopped, the solvent was removed and further purification was conducted by using alumina column chromatography to get crimson microcrystal. Yield, 51.3%, 49.7 mg. Anal. Calcd. for C_48_H_35_N_7_SC1Ir (%): C, 59.46; H, 3.64; N, 10.11. Found (%): C, 59.19; H, 3.91; N, 10.01. ^1^H NMR (500 MHz, CD_3_OD): δ 8.78 (d, *J* = 1.5 Hz, 1H), 8.57 – 8.48 (m, 3H), 8.30 (s, 3H), 8.27 (dd, *J* = 6.0, 1.5 Hz, 1H), 8.23 (d, *J* = 6.0 Hz, 1H), 8.08 (t, *J* = 8.0 Hz, 2H), 7.87 (t, *J* = 8.0 Hz, 2H), 7.59 (s, 1H), 7.57 – 7.51 (m, 4H), 7.42 (t, *J* = 10.0 Hz, 2H), 7.32 – 7.26 (m, 2H), 7.11 (d, *J* = 12.0 Hz, 2H), 6.89 (d, *J* = 8.0 Hz, 2H), 6.72 (dd, *J* = 18.0, 7.5 Hz, 2H), 3.40 (d, *J* = 15.1 Hz, 6H), 2.49 (s, 3H). ^13^C NMR (125 MHz, CD3OD) δ164.62, 164.55, 162.30, 156.57, 154.36, 153.18, 152.99, 152.87, 152.81, 152.68, 148.37, 146.82, 144.41, 143.47, 140.06, 139.90, 139.85, 130.62, 130.59, 129.97, 129.88, 129.80, 129.17, 128.66, 127.15, 127.01, 125.14, 125.01, 123.74, 122.98, 122.02, 120.63, 26.14, 26.09, 19.79. ES-MS (CH3OH): m/z = 933.95 [M-Cl^-^]^+^.

^1^H and ^13^C NMR spectra were recorded using a Bruker 500 Nuclear Magnetic Resonance Spectrometer using CDCl_3_ or CD_3_OD as the deuterated solvent. The electronic absorption spectra were recorded using a Perkin-Elmer Lambda 850 UV/Vis spectrometer. The emission spectra were recorded using a Perkin-Elmer LS 55 luminescence spectrometer and FLS 980 luminescence spectrometer. Microanalysis (C, H, and N) was carried out using an Elemental Vario EL elemental analyzer. Electro spray mass spectra were recorded using an LCQ system (Finnigan MAT, USA).

### Cell culture

HeLa cells were gifted from Dr. Carolyn M. Price lab (University of Cincinnati). Penta knockout HeLa cells were gifted from Dr. Richard J. Youle lab (National Institutes of Health). Cells were cultured in Dulbecco’s modified Eagle medium supplemented with 10% FBS, penicillin (100 units/ml), and streptomycin (100 μg/ml) in a 5% CO_2_ humidified incubator at 37 °C.

### Cell viability and cytotoxicity assay

Cells were treated in a 96-well plate at density of 5 × 10^5^ cell/ml. The viability was determined by using a cell counting kit-8 (CCK-8). 10 μl CCK-8 solution was added to each well and the OD value for each well was read at wavelength 450 nm on a microplate reader (Multiskan, Thermo, USA).

### Flow cytometry analysis

Cells were seeded on 6-well plate at density of 1 × 10^5^ cell/ml in 1 ml of complete medium for 24 h. After treatment with iridium(III) complex dye (0.1, 0.25, 0.5, 1, 2.5, 5, 10, 25, and 50 μM) for 30 min, cells were collected by trypsinization and washed 2 times with cool PBS. Cells were resuspended by 500 μl binding buffer while avoiding light prior to detection by flow cytometry.

### Live cell labeling

Cells were incubated with 1 μM IR for 30 min in free DMEM, and washed with free DMEM 3 times and observed using a fluorescence microscope (CX41-32RFL; Olympus, Japan), confocal laser scanning microscopy or OMX 3D-SIM super-resolution microscope.

### Confocal laser scanning microscopy

The images were obtained using a LSM-710 confocal laser scanning microscope (Carl Zeiss, Inc.) equipped with a 63×/1.49 numerical aperture oil-immersion objective lens and were analyzed with ZEN 2012 (Carl Zeiss, Inc.) and ImageJ software (National Institutes of Health). All fluorescence images were analyzed and the background subtracted with ImageJ software. Pearson’s coefficient was quantified using the Colocalization Analysis plugin for ImageJ.

### OMX 3D-SIM super-resolution microscope imaging

Super-resolution images were acquired on OMX 3D-SIM Microscope (Bioptechs, Inc) equipped with a Olympus 100×/1.49 numerical aperture oil-immersion objective lens and solid-state lasers. Images were captured with an electron-multiplying charge coupled device (EMCCD) camera (Photometrics Cascade II) with a gain value of 3000 at 10 MHz. The exposure time was set to 50 ms for each raw data capture. Picture was obtained at 512 × 512 using Z-stacks with step size of 0.2 μm. SIM frames were deliberately spaced at 1-s, 2-s, 8-s or 15-s intervals according to the purpose of each experiment. SIM images were analyzed with Nikon Elements and ImageJ software.

## Acknowledgments

This research was supported by 973 Program (Nos. 2015CB856301 and 2015CB856304) from Ministry of Science and Technology of China, the National Science Foundation of China (Nos. 21525105, 21471164, 21778079 and 21701196), the Fundamental Research Funds for the Central Universities (7lgjc11), China Postdoctoral Science Foundation (20173100041090767), Natural Science Foundation of Shandong Province (ZR2017PH072), Key Research and Development Plan of Shandong Province(2018GSF 121033), and National Institutes of Health (NIH, R01NS094144 and R01CA211066 to J.G.). The Light Microscopy Imaging Center (LMIC) is supported in part with funds from Indiana University Office of the Vice Provost for Research. The OMX 3D-SIM microscope was provided by NIH grant S10 RR028697. We also thank Dr. Ehmer Birgit at University of Cincinnati for assistance with laser scanning confocal microscope and flow cytometry.

## Author contributions

C.J., R.G. and H.C. designed, synthesized and characterized iridium(III) complex dye. Q.C. and Z.T. collected all OMX 3D-SIM super-resolution microscope data. Q.C and X.S. analyzed and processed the OMX 3D-SIM data. Q.C., X.S., and C.W. cultured cell and performed confocal laser scanning microscopy. F.L. performed cytotoxicity assay and flow cytometry analysis. J.G., L.J., F.W., H.C., and J.D. conceived the project, designed the experiments, and wrote the manuscript with the help of all authors.

